# The generality of post-antimicrobial treatment persistence of replicatively-attenuated *Borrelia burgdorferi* in a mouse model

**DOI:** 10.1101/596122

**Authors:** Emir Hodzic, Denise M. Imai, Edlin Escobar

**Affiliations:** Real-Time PCR Research and Diagnostic Core Facility, School of Veterinary Medicine, University of California at Davis, One Shields Avenue, California, United States of America; Comparative Pathology Laboratory, School of Veterinary Medicine, University of California at Davis, One Shields Avenue, California, United States of America

## Abstract

A basic feature of infection caused by *Borrelia burgdorferi*, the etiological agent of Lyme borreliosis, is that persistent infection is the rule, not the norm, in its many hosts. The ability to persist and evade host immune clearance poses a challenge to effective antimicrobial treatment. A link between therapy failure and the presence of persister cells has started to emerge. There is growing experimental evidence that viable, but non-cultivable spirochetes persist following treatment with several different antimicrobial agents, then resurge after 12 months. The current study utilized the mouse model to evaluate if persistence and resurgence occur following antimicrobial treatment in a disease-susceptible (C3H/HeN) and disease-resistant (C57BL/6) mouse strain infected with *B. burgdorferi* strains N40 and B31, to confirm the generality of these phenomena. The status of infection was evaluated at 12 and 18-months after treatment. The results demonstrated that persistent spirochetes remain viable for up to 18 months following treatment, but divide slowly, thereby being tolerant to the effects of antimicrobial agents, as well as being non-cultivable. The phenomenon of persistence and resurgence in disease-susceptible C3H mice is equally evident in disease-resistant B6 mice, and not unique to any particular *B. burgdorferi* strain. The results also demonstrate that following antimicrobial treatment, both strains of *B. burgdorferi*, N40 and B31, lose one or more small plasmids, resulting in attenuation. The biological relevance of attenuated *B. burgdorferi* spirochetes is probably inconsequential. The study demonstrated that non-cultivable spirochetes can persist in a host following antimicrobial treatment for a long time but did not demonstrate their clinical relevance in a mouse model of chronic infection.

## INTRODUCTION

*Borrelia burgdorferi*, the etiological agent of Lyme borreliosis has become an important public health problem and the most prevalent tick-borne disease in the United States (1, 2). It occurs across much of the northern hemisphere, causing considerable morbidity and in some cases mortality in humans, domestic animals, and occasionally wildlife. It is the most frequently diagnosed tick-borne disease in the United States. According to the Centers for Disease Control and Prevention (CDC), it is estimated there are more than 300,000 human cases of Lyme borreliosis annually (3), and is expecting to rise (4).

Untreated human Lyme borreliosis results in disease with a wide range of clinical symptoms and protean manifestation, depending on the stage of infection (5-9). The treatment with antimicrobial agents of patients with diagnosed Lyme borreliosis mainly resolves the objective clinical manifestations. Associated subjective symptoms often may persist for many weeks, even months (10). However, a proportion of patients remain ill (11, 12) and delayed treatment is associated with negative clinical outcomes (13, 14). There are many reports that despite resolution of the clinical signs of infection after treatment with antimicrobial agents, a minority of patients experience chronic Lyme disease (15-22).

A hallmark of Lyme borreliosis is persistent infection. This has been experimentally proven in *Peromyscus* mice (23), laboratory mice (24), rats (25), hamsters (26), gerbils (27), guinea pigs (28), dogs (29), and non-human primates (30), and in reported confirmed spontaneous cases in humans based on culture (31-37) and PCR (38-41). Chronic cases require prolonged treatment, and treatment is often less effective (15, 42-44). Several reports have provided evidence of *B. burgdorferi* presence in collagenous tissues, following antimicrobial therapy during chronic infection in different animal models, including mice (45-50), dogs (29, 51, 52), and non-human primates (48, 53, 54), and in spontaneous cases in humans (17-21). What is unique about all of these studies is that spirochetes can be detected by PCR for *B. burgdorferi*-specific DNA (BbDNA), but not by culture. There is also strong scientific evidence that uninfected ticks were able to acquire *B. burgdorferi* that survive various antimicrobial treatments and transmit them to naïve hosts following the molt to the next stage (45-47, 50, 54).

The nature of the formation of persistent spirochetes is still unknown and is matter of speculation. Several mechanisms have been proposed that involve a suppression of the innate and adaptive immune systems, complement inhibition, induction of anti-inflammatory cytokines, formation of immune complexes, antigenic variation, and physical seclusion (55-59). *B. burgdorferi* has evolved to persist in immunologically competent hosts as a survival strategy for maintaining its natural host-vector life cycle (27, 53, 55, 60). Persistence of non-cultivable spirochetes has been shown to occur following treatment with several different classes of antimicrobial agents, and the phenomenon is likely explained by antimicrobial tolerance (in contrast to antimicrobial resistance or inadequate antimicrobial treatment), in which all classes of antimicrobials fail to completely eliminate non-dividing or slowly-dividing subpopulations of a broad array of bacteria and fungi (61, 62). It has been known for decades that during in vitro passage, *B. burgdorferi* is highly prone to plasmid loss (63-65), and therefore plasmid loss is likely to also occur during the course of infection and increase over time.

In mouse studies performed in this laboratory, mice were treated with ceftriaxone, doxycycline, or tigecycline at various intervals of infection, and tissues were tested at intervals after treatment. Tissues remained BbDNA PCR-positive up to 12 months but were consistently culture-negative. Morphologically intact spirochetes could be visualized by immunohistochemistry in tissues from treated mice; ticks could acquire morphologically intact *B. burgdorferi* and BbDNA from treated mice; ticks remained BbDNA-positive through molting into nymphs and adults; nymphs transmitted BbDNA to recipient immunocompromised mice; allografts from treated mice transplanted into recipient immunocompromised mice transferred BbDNA to recipient mice; and both tick- and transplant-inoculated mice had disseminated BbDNA. BbDNA-positive tissues were also positive for *B. burgdorferi*-specific RNA transcription. Furthermore, quantitative PCR indicated low-levels of replication during these various stages (46, 47, 50). In the most recent study we observed the resurgence of non-cultivable spirochetes in assessed tissues of antimicrobial treated mice after 12 months, and the overall tissue spirochete burden reached the levels detected in sham-treated mice at the same time point. Despite the continued non-cultivable state, RNA transcription of multiple *B. burgdorferi* genes was detected in host tissues, BbDNA was acquired by xenodiagnostic ticks, and spirochetal forms could be visualized within ticks and mouse tissues by immunofluorescence and immunohistochemistry, respectively. A number of host cytokines were up- or down-regulated in tissues of both saline- and antimicrobial-treated mice in the absence of histopathology, indicating host response to the presence of non-cultivable spirochetes, despite the lack of inflammation in tissues (47).

We hypothesize that during the course of infection, *B. burgdorferi* proliferates and incidentally generates an increasingly heterogeneous population of replicatively-attenuated spirochetes that have lost one or more small plasmids. These “attenuated” spirochetes remain viable, but because of their plasmid loss, they divide slowly, thereby being tolerant to the effects of antimicrobials, as well as being non-cultivable. Results obtained from this study demonstrated the generality of spirochete persistence and in particular, resurgence, following antimicrobial treatment by demonstrating the phenomena in genetically susceptible C3H/HeN (C3H) and resistant C57BL/6 (B6) mouse strains infected with *B. burgdorferi* strain N40 compared to strain B31 for up to 18 months after treatment.

## MATERIAL AND METHODS

### Mouse infections

All experiments involving vertebrate animals were approved by the Institutional Animal Care and Use Committee (IACUC) at University of California Davis (UCD). All animals were purchased from The Jackson Laboratory, Bar Harbor, Maine, and were cared for by staff from Teaching and Research Animal Care Services (TRACS), at UCD. Mice were maintained in an isolated room within filter-top cages and were provided food and water *ad libitum*. The animals used in this study were C3H and B6 specific-pathogen-free 5-week-old female mice and were maintained in cohorts of 5 mice per cage under identical husbandry conditions in the same animal room. Euthanasia was performed by rapid carbon dioxide narcosis in accordance with guidelines of the American Veterinary Medical Association (AVMA) (66). C3H and B6 mice were randomly divided into two equal groups. One group of C3H and B6 mice was infected with *B. burgdorferi* N40 and another one with *B. burgdorferi* B31. Each mouse was infected by syringe inoculation of 10^4^ *B. burgdorferi* spirochetes at the mid-log phase in ml of Barbour-Stoenner-Kelly-II (BSK-II) medium intradermally at the dorsal thoracic midline.

### B. burgdorferi culturing

Two low-passage clonal strains of *B. burgdorferi* sensu stricto were used. *B. burgdorferi* strains N40 and B31 were cloned by threefold limiting dilution *in vitro* and passage in mice to prove infectivity and pathogenicity as described previously (24). Frozen aliquots of clonal strains N40 and B31 were thawed and cultured in modified BSK-II medium (67) at 33°C. Spirochetes were assessed for viability and enumerated by dark-field microscopy using a Petroff-Hausser bacterial counting chamber (Hausser Scientific, Horsham, PA) immediately prior to use, and diluted to appropriate concentrations in BSKII medium. Mice were inoculated subdermally on the dorsal thoracic midline with 10^4^ mid-log phase spirochetes in 0.1 ml of BSK-II medium. Infection status with cultivable *B. burgdorferi* was determined by culture of urinary bladder and sub-inoculation site (deep dermis). Tissues were collected from mice aseptically at necropsy, and then cultured in medium without antibiotic, as described (68).

To determine if resurgent spirochetes regain cultivability at 18 months after treatment and to increase their detectability, multiple tissue sites were collected and cultured in four different media. In addition to urinary bladder and inoculation site we also collected front joint, ear, quadriceps muscle, and spleen. Heart tissue was not available for culture as was used for PCR and histology. To facilitate cultivation, several media have been introduced, such as BSK-II medium, modified Kelly-Pettenkofer (MKP) medium, BSK-II + agarose, and BSK-II + carbohydrates medium (cordially provided by Monica E. Embers of Tulane National Primate Research Center, Covington, Louisiana). However, these media differ in their potential to support the growth of borreliae.

### Xenodiagnosis

Laboratory-reared, pathogen-free *Ixodes scapularis* larvae were provided by Melissa J. Caimano of Department of Molecular Biology and Biophysics, UConn Health, Farmington, CT. All larvae were derived from single gravid ticks for each study. Around 40 larval ticks were placed on each mouse 1 week prior to necropsy, allowed to feed to repletion, collected, and then allowed to molt and harden into nymphs. Cohorts of ticks collected from each mouse were maintained separately, so that ticks within each cohort could be tested individually or in groups by qPCR. Nymphal ticks were frozen in liquid nitrogen, ground with a mortar and pestle, and DNA was extracted for PCR analysis.

### Antimicrobial treatment and monitoring

The treatment with ceftriaxone was initiated during the early immune/dissemination stage at 4 weeks post infection of *B. burgdorferi* in immunocompetent C3H and B6 mice to mimic the stages found in human disease. Previous studies in this laboratory have shown that the minimal inhibitory concentration (MIC) and minimal bactericidal concentration (MBC) of ceftriaxone is 0.015 μg/ml and 0.06 μg/ml, respectively. The experimental design for this experiment involved eight treatment groups, as depicted in Table 1. A ceftriaxone dosage of 16 mg/kg in 500 μl 0.9% normal saline was administrated intraperitoneally in mice twice daily for 5 days, followed by once daily for 23 days. Over a 4-h period, serum concentrations of ceftriaxone in mice were measured on the day of treatment by bleeding mice at 0, 2 and 4 hours after treatment. Average serum concentrations of ceftriaxone were 18.625 μg/ml, and 1.795 μg/ml at 2 and 4 hours, respectively. As expected, intraperitoneally administrated ceftriaxone accumulated rapidly in the circulation and then decreased gradually during the following 2 hours. Studies with ceftriaxone in mice have been challenged because the pharmacodynamics (T>MIC) of ceftriaxone do not parallel those in humans. However, the ceftriaxone treatment regimen used in our studies transiently achieves more than adequate serum concentrations. No differences were observed in serum ceftriaxone concentrations between different mouse strains (C3H and B6).

**Table 1.** The experimental design for the executed experiment involved eight treatment groups.

| Group | *B. burgdorferi* | Mouse strain | Treatment | 12 months | 18 months | n= |
| --- | --- | --- | --- | --- | --- | --- |
| 1 | N40 | C3H | ceftriaxone | 5 | 5 | 10 |
| 2 | B31 | C3H | ceftriaxone | 5 | 5 | 10 |
| 3 | N40 | B6 | ceftriaxone | 5 | 5 | 10 |
| 4 | B31 | B6 | ceftriaxone | 5 | 5 | 10 |
| 5 | N40 | C3H | saline | 5 | 5 | 10 |
| 6 | B31 | C3H | saline | 5 | 5 | 10 |
| 7 | N40 | B6 | saline | 5 | 5 | 10 |
| 8 | B31 | B6 | saline | 5 | 5 | 10 |

### Tissue processing for molecular analysis

Mouse tissue samples for molecular analysis were processed as previously described (47, 69), with slight modifications. Briefly, samples of the heart base, ventricular myocardium (the heart was bisected through the atria and ventricles, with one-half of the heart base and ventricles used for nucleic acid extraction and the other half processed for histology), and the left tibiotarsal joint were collected from each mouse. The samples were collected aseptically, immediately weighed, snap-frozen in liquid nitrogen, and stored at −80°C before nucleic acid extraction. Total nucleic acid was extracted with QIAcube HT system (Qiagen, Valencia, Calif.), according to the manufacturer’s instructions for tissue or insects. The system enables semi-automated high-throughput nucleic acid purification, yielding high-quality pathogen nucleic acids free of contaminants and inhibitors. The copy number of each *B. burgdorferi* target gene was expressed per milligram of tissue weight.

### Molecular quantitative analysis

Because low DNA/RNA copy numbers were expected in tissues from antimicrobial-treated mice, each sample was subjected to pre-amplification. We used the same *B. burgdorferi* outer surface protein (*ospA*) and flagellin (*flaB*), and a mouse glyceraldehyde-3-phosphate dehydrogenase (*gapDH*) assay primers, standardized and optimized for real-time quantitative PCR (qPCR), as described (70). Purified DNA/buffer was heated for 5 minutes at 95°C then placed on ice. Each reaction contained Advantage buffer, Advantage 2 Polymerase (Takara Bio, Mountain View, CA), mM each dNTP, 10 μM of each primer, and DNA template (100 ng/μl). Standard amplification conditions were as follows: 1 min at 94°C, 25 cycles of 15 s at 94°C, 15 s at 55°C, and 45 s at 70°C, 5 min at 70°C, and then 10 min at 4°C. The pre-amplified products were diluted at a ratio of 1∶10 and used as templates for qPCR analysis. To assess transcriptional activity of *B. burgdorferi flaB* and *16S rRNA* genes, first-strand cDNA was synthesized from total RNA using the QuantiTect Reverse Transcription Kit (Qiagen) in 50-μL reactions, as described (70). To increase fidelity, efficiency, and greater yield of cDNA, the Advantage® 2 Polymerase Mix (Clontech Laboratories) was used, according to the manufacturer’s instructions.

Two specific primers and one internal, fluorescence-labeled probe were designed with Primer Express^TM^ software (ThermoFisher Scientific), as described (71). The amplification efficiency (E) of all assays was calculated from the slope of a standard curve generated on a 10-fold dilution in triplicate for every DNA sample, using the formula E = 10^(−1/slope)^ −1. In order to obtain accurate and reproducible results, all assays were determined to have an efficiency of >95%. To quantify the copy numbers of DNA target genes, plasmid standards were prepared in order to create absolute standard curves, as described (71). Based on the amplification efficiencies, detection limits were approximately 10 copies of DNA per reaction, and the analytical sensitivity for each target gene was in the range from <10 to 10^9^ copies. All samples were assayed in duplicate with positive and negative controls by the qPCR optimized assays and analyzed for the presence of mouse *gapDH* in order to determine the efficiency of the nucleic acid extraction, amplification, and as an indicator of inhibition. Amplification, data acquisition, and data analysis were performed on a 7900HT Fast Real-Time PCR System (ThermoFisher, Scientific). The thermal cycling conditions were as follow: 2 min at 50°C to 10 min at 94.5°C, followed by 40 cycles of denaturation at 97°C for 30 sec, and annealing and extension at 59.7°C for 1 min.

### Histology

Formalin-fixed rear legs (demineralized in 15% formic acid) and hearts were routinely processed for histology, paraffin-embedded and sectioned at 5 μm. Sections were stained with hematoxylin and eosin. All tissues were evaluated by a board-certified veterinary anatomic pathologist. Legs were examined for evidence of arthritis and synovitis involving the knee and tibiotarsal joints as characterized by examination of infiltration of neutrophils, synovial proliferation, and exudation of fibrin into joint or tendon sheath lumina as well as neutrophil and macrophage infiltration of tissues at the base of the heart (myocarditis). All determinations were made in a blinded fashion.

### Statistics

Statistical analysis of qPCR data between antimicrobial treated and sham treated mice at different time points were performed using independent samples *t*-test or one-way analysis of variance, followed by multiple pairwise comparisons by Tukey’s honestly significant difference (HSD) test (SPSS 16.0 for Mac; SPSS Inc., Chicago, IL). Differences were considered significant with p≤0.5.

To statistically evaluate the severity of inflammation in collected tissue between antimicrobial treated and sham treated mice Kruskal-Wallis test was performed using GraphPad Prism, 7.03.

## RESULTS

### Persistence and resurgence of *B. burgdorferi* N40 and B31 following antimicrobial treatment in C3H and B6 mice

Mouse tissue samples, heart base, ventricular muscle, and tibiotarsal joint, were collected aseptically, immediately weighed, snap-frozen in liquid nitrogen, and stored at −80°C before total nucleic acid extraction. Quality control of qPCR was assessed by targeting mouse *gapDH* gene. Spirochetal DNA was *flaB* and *ospA*. Quantitative data were expressed as the number of DNA copies per milligram of tissue weight. Mouse *gapDH* qPCR was positive from all assessed samples (average Cq ± SD, 19.36 ± 4.69), confirming successful extraction of total nucleic acid.

Based upon *flaB*/*ospA* DNA qPCR on assessed tissue samples, all C3H and B6 mice inoculated with either *B. burgdorferi* strains N40 or B31 treated with saline were qPCR-positive at 12 and 18 months after treatment in most tissues. The data in Table 2 and Table 3 summarize positive or negative *flaB/ospA* DNA qPCR results. Among ceftriaxone-treated C3H and B6 mice infected with N40 or B31, 100% and 74% tested positive for *flab/ospA* DNA qPCR at tibiotarsal joints at 12 and 18 months after treatment, respectively. Sporadically positive heart base and ventricular muscle samples were detected in mice at 12 months after antimicrobial treatment (Table 2), but not in mice at 18 months (Table 3).

**Table 2.**
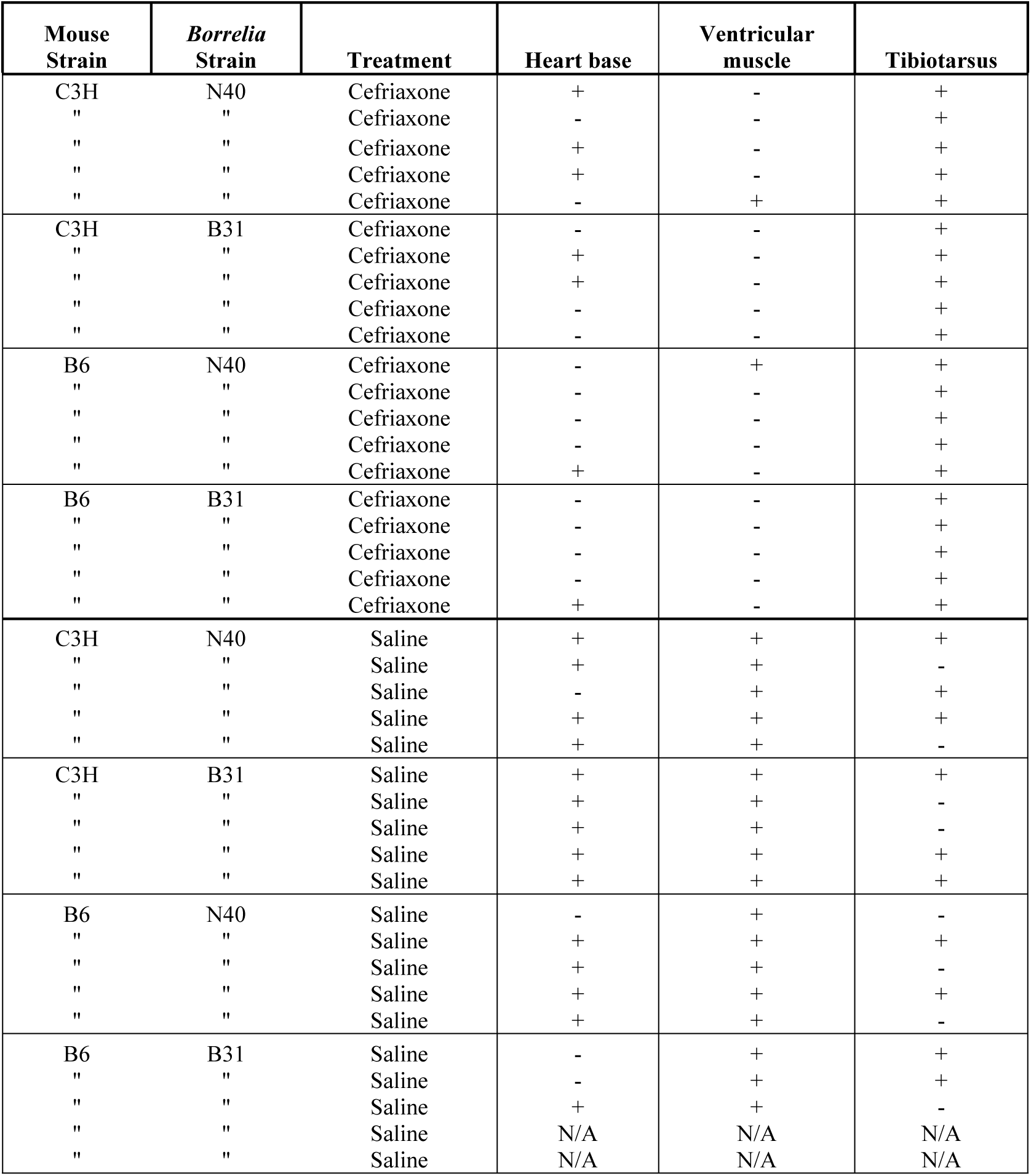
Analysis of *flaB/ospA* DNA in heart base, ventricular muscle, and tibiotarsus, from *B. burgdorferi* infected mice treated with ceftriaxone and saline solution, then necropsied **12 months** after completion of treatment

**Table 3.** Analysis of *flaB/ospA* DNA in heart base, ventricular muscle, and tibiotarsus, from *B. burgdorferi* infected mice treated with ceftriaxone and saline solution, then necropsied **18 months** after completion of treatment

| Mouse Strain | *Borrelia* Strain | Treatment | Heart base | Ventricular muscle | Tibiotarsus |
| --- | --- | --- | --- | --- | --- |
| C3H | N40 | Ceftriaxone | - | - | - |
| " | " | Ceftriaxone | - | - | - |
| " | " | Ceftriaxone | - | - | + |
| " | " | Ceftriaxone | - | - | - |
| " | " | Ceftriaxone | N/A | N/A | N/A |
| C3H | B31 | Ceftriaxone | - | - | + |
| " | " | Ceftriaxone | - | - | + |
| " | " | Ceftriaxone | - | - | + |
| " | " | Ceftriaxone | - | - | + |
| " | " | Ceftriaxone | - | - | + |
| B6 | N40 | Ceftriaxone | - | - | - |
| " | " | Ceftriaxone | - | - | + |
| " | " | Ceftriaxone | - | - | + |
| " | " | Ceftriaxone | - | - | + |
| " | " | Ceftriaxone | - | - | + |
| B6 | B31 | Ceftriaxone | - | - | + |
| " | " | Ceftriaxone | - | - | + |
| " | " | Ceftriaxone | - | - | + |
| " | " | Ceftriaxone | - | - | + |
| " | " | Ceftriaxone | - | - | + |
| C3H | N40 | Saline | + | + | + |
| " | " | Saline | + | + | + |
| " | " | Saline | - | + | + |
| " | " | Saline | + | + | + |
| " | " | Saline | N/A | N/A | N/A |
| C3H | B31 | Saline | + | + | + |
| " | " | Saline | + | + | + |
| " | " | Saline | + | + | + |
| " | " | Saline | N/A | N/A | N/A |
| " | " | Saline | N/A | N/A | N/A |
| B6 | N40 | Saline | - | + | + |
| " | " | Saline | + | + | + |
| " | " | Saline | + | + | + |
| " | " | Saline | + | + | + |
| " | " | Saline | + | + | + |
| B6 | B31 | Saline | + | + | + |
| " | " | Saline | + | + | + |
| " | " | Saline | + | + | + |
| " | " | Saline | + | + | + |
| " | " | Saline | + | + | + |

Quantification of each target gene in assessed samples was accomplished by preparing plasmid standards in order to create absolute standard curves to determine starting copy number. Mean DNA copy numbers of *flaB/ospA* genes in collected tissue samples among different mouse strains (C3H and B6) and different *B. burgdorferi* stra ins (N40 and B31) of ceftriaxone-treated groups at 12 months after treatment were nearly equivalent (mean ± SD, 2.97 x 10^1^ ± 1.72 x 10^1^). Among saline-treated groups significantly higher DNA copy numbers of *flaB/ospA* were determined in C3H mice infected with N40 and B31 in comparison to B6 mice infected with N40 and B31 (*P*<0.05) (Fig 1). Significantly higher copy numbers of *flaB/ospA* were determined in tissue samples of saline-treated mice in comparison to the ceftriaxone-treated mice (*P*<0.05) (Fig 1). Thus, these results suggested resurgence of spirochetes at 12 months following antimicrobial treatment in both C3H and B6 mice infected with both *B. burgdorferi* strains, N40 and B31, although all mice remained culture-negative. qPCR results also revealed that a population of resurged spirochetes that survived antimicrobial treatment and detected at 12 months, remained in mice even after 18 months. They were exclusively detected in tibiotarsal joints of all treatment groups, with mean copy numbers of *flaB/ospA* per milligram tissue that were nearly equivalent (mean ± SD, 1.52 x 10^1^ ± 9.74 x 10^0^) (Fig 2). qPCR results of saline treated groups at 18 months were similar to the results of the same treated groups at 12 months. *flaB/ospA* genes copies among saline treated groups at 18 months were lower in B6 mice infected with N40 and B31 in comparison to C3H mice infected with N40 and B31, but not significantly. Mean *flaB/ospA* copy numbers in tibiotarsal joints at 18 months were significantly higher (*P*<0.05) in saline treated C3H mice infected with N40 and B31 in comparison to the same groups of antimicrobial treated mice (Fig 2).

**Figure 1.**
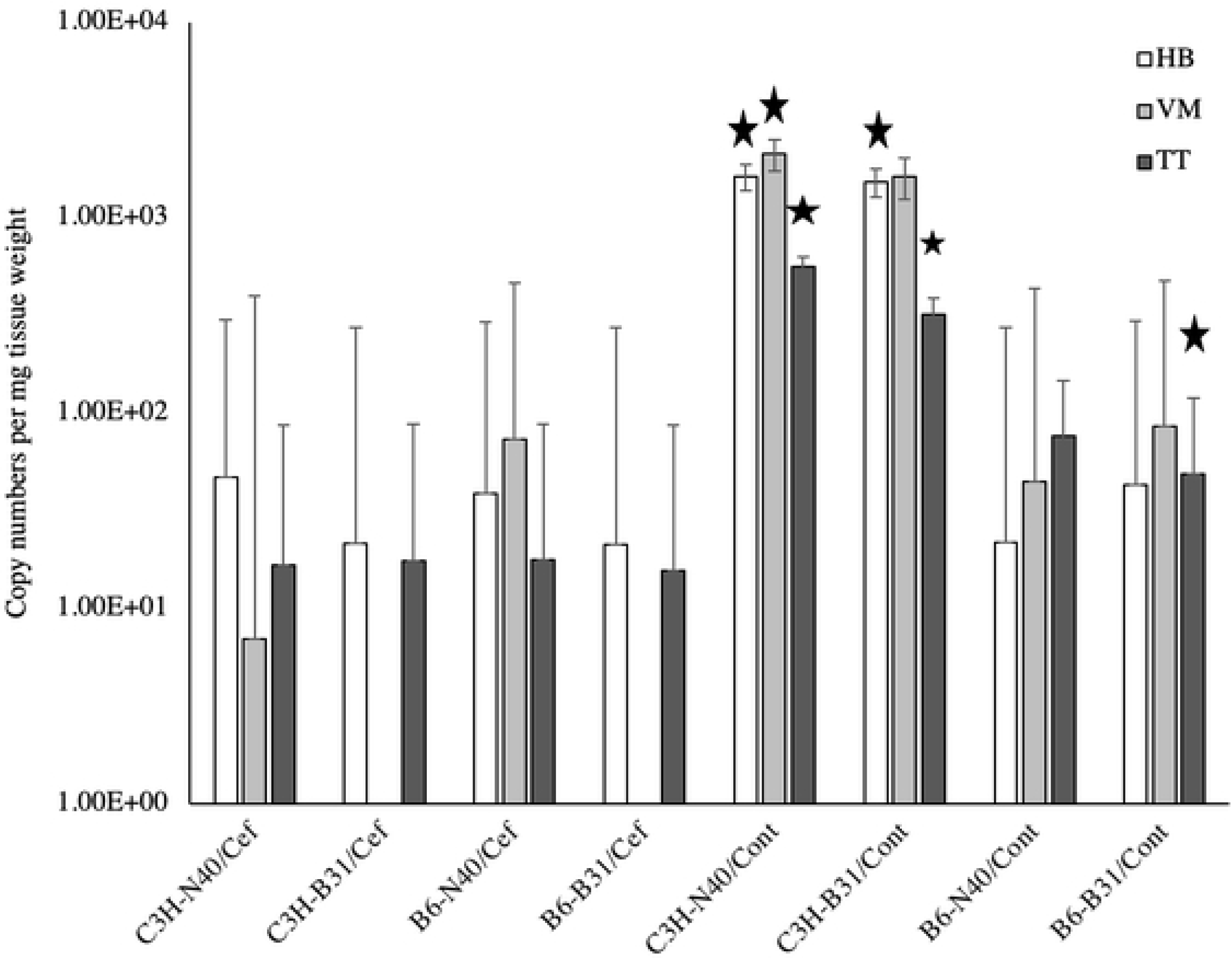
Copy numbers of *B. burgdorferi* DNA of *flaB/ospA* genes in heart base (HB), ventricular muscle (VM), and tibiotarsus (TT) of C3H and B6 mice infected with either N40 or B31 strains at 12 months after ceftriaxone and saline solution treatment. *Significantly higher copy number (p<0.05)

**Figure 2.**
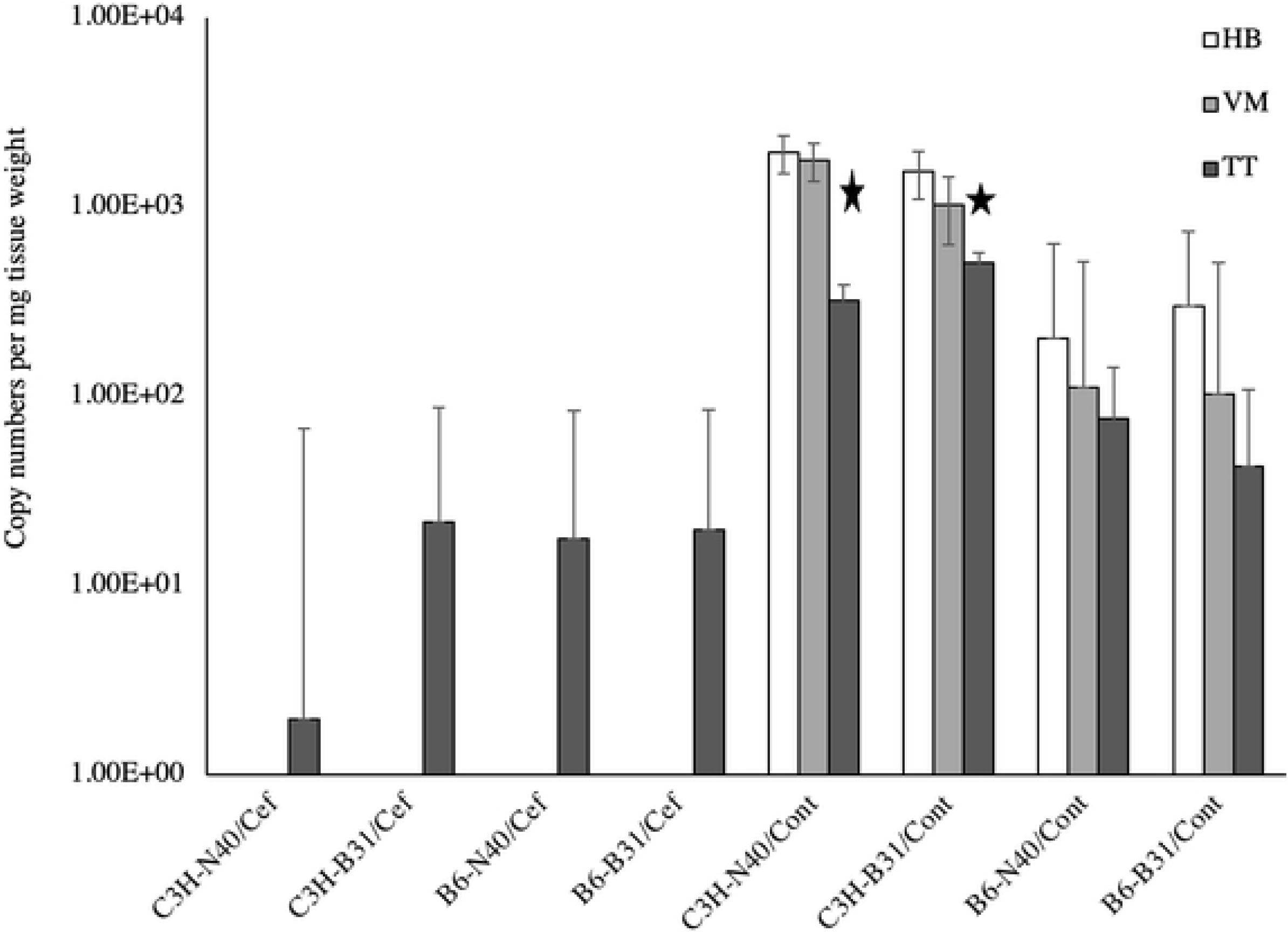
Copy numbers of *B. burgdorferi* DNA of *flaB/ospA* genes in heart base (HB), ventricular muscle (VM), and tibiotarsus (TT) of C3H and B6 mice infected with either N40 or B31 strains at 18 months after ceftriaxone and saline solution treatment. *Significantly higher copy number (p<0.05)

Analysis of *B. burgdorferi flaB* and *16S rRNA* transcriptional activity revealed that most of the saline treated C3H and B6 mice infected with N40 or B31 strains were positive. Transcriptional activity of *flaB* in samples from antimicrobial treated mice (C3H and B6) necropsied at 12 and 18 months after treatment were 52% and 35%, respectively, while *16S rRNA* was constitutively detected in all samples. The results indicate viability among the persisting spirochetes.

### Confirmation of spirochetes non-cultivability at 12 and 18 months following antimicrobial treatment

The infection status of mice at 12 months after antimicrobial treatment was assessed by culture of inoculation site and urinary bladder. For culture, we used a modified BSK-II medium. The sensitivity of BSK-II medium for detection of viable spirochetes was verified by serial 10-fold dilutions of a *B. burgdorferi* N40 and B31 culture, as described (46). The culture results revealed that all saline treated mice were positive, with significant numbers of spirochetes detected (Table 4). When mice treated with ceftriaxone evaluated at 12 months after completion of treatment, none of the tissues were culture positive. In single culture two non-motile spirochetes were observed (Table 4). Resurgence at 12 months after treatment suggests that spirochetes have regained some semblance of replicative activity and/or are disseminating once again within mouse tissues.

**Table 4.** Culture results for individual mice treated with ceftriaxone and saline solution commencing at 4 weeks after inoculation, then subjected to necropsy at 12 months after completion of treatment

| Mouse Strain | *Borrelia* Strain | Treatment | Inoculation site | Urinary bladder |
| --- | --- | --- | --- | --- |
| C3H | N40 | Ceftriaxone | - | - |
| " | " | Ceftriaxone | - | - |
| " | " | Ceftriaxone | - | - |
| " | " | Ceftriaxone | - | - |
| " | " | Ceftriaxone | - | - |
| C3H | B31 | Ceftriaxone | N/A | N/A |
| " | " | Ceftriaxone | - | - |
| " | " | Ceftriaxone | - | - |
| " | " | Ceftriaxone | - | - |
| " | " | Ceftriaxone | - | - |
| B6 | N40 | Ceftriaxone | - | - |
| " | " | Ceftriaxone | - | - |
| " | " | Ceftriaxone | - | - |
| " | " | Ceftriaxone | - | - |
| " | " | Ceftriaxone | - | + |
| B6 | B31 | Ceftriaxone | - | - |
| " | " | Ceftriaxone | - | - |
| " | " | Ceftriaxone | - | - |
| " | " | Ceftriaxone | - | - |
| " | " | Ceftriaxone | - | - |
| C3H | N40 | Saline | ++ | +++ |
| " | " | Saline | ++ | +++ |
| " | " | Saline | ++ | +++ |
| " | " | Saline | + | +++ |
| " | " | Saline | + | ++ |
| C3H | B31 | Saline | ++ | +++ |
| " | " | Saline | + | + |
| " | " | Saline | + | ++ |
| " | " | Saline | ++ | ++ |
| " | " | Saline | ++ | ++ |
| B6 | N40 | Saline | ++ | + |
| " | " | Saline | + | + |
| " | " | Saline | +++ | + |
| " | " | Saline | ++ | + |
| " | " | Saline | ++ | +++ |
| B6 | B31 | Saline | +++ | +++ |
| " | " | Saline | + | + |
| " | " | Saline | ++ | ++ |
| " | " | Saline | N/A | N/A |
| " | " | Saline | N/A | N/A |

For the first time point of this study (12 months), a limited number of sample sites (urinary bladder and inoculation site) were cultured. These are not optimal sites for culture after antimicrobial treatment, since persisting spirochetes tend to be localized to heart base and joint tissue. To determine if resurgent spirochetes regain cultivability at 18 months after treatment, multiple sample types were collected such as joint, ear, quadriceps muscle, spleen, urinary bladder, and inoculation site. Heart tissue was not available for culture as was used for PCR and histology. To facilitate cultivation, several media have been introduced, such as BSK-II medium, modified Kelly-Pettenkofer (MKP) medium, BSK-II + agarose, and BSK-II + carbohydrates medium. The inoculated cultures were incubated for 50 days at 33°C and examined weekly. Culture tubes were gently but thoroughly stirred, and a drop of culture was examined by dark-field microscopy. The culture results revealed that all saline treated mice were positive, with significant numbers of motile spirochetes detected (Table 5, Fig 3A) in the majority of tissue samples cultured. When mice treated with ceftriaxone were evaluated, one ear tissues sample from B6 mouse infected with B31 (Fig 3B) and one front joint tissue sample from B6 mouse infected with N40 (Fig 3C) were culture positive (Table 5). Tissues were cultured in MKP and BSK-II + carb medium, respectively. After 4 weeks of incubation spirochetes in cultured medium of ceftriaxone treated mice appeared non-motile with irregular curves (dead), so we were unable to subculture them. These results confirm our previous findings that antimicrobial-treated spirochetes are not cultivable, probably as a result of being genetically impaired (plasmid loss) after treatment with antimicrobials.

**Table 5.**
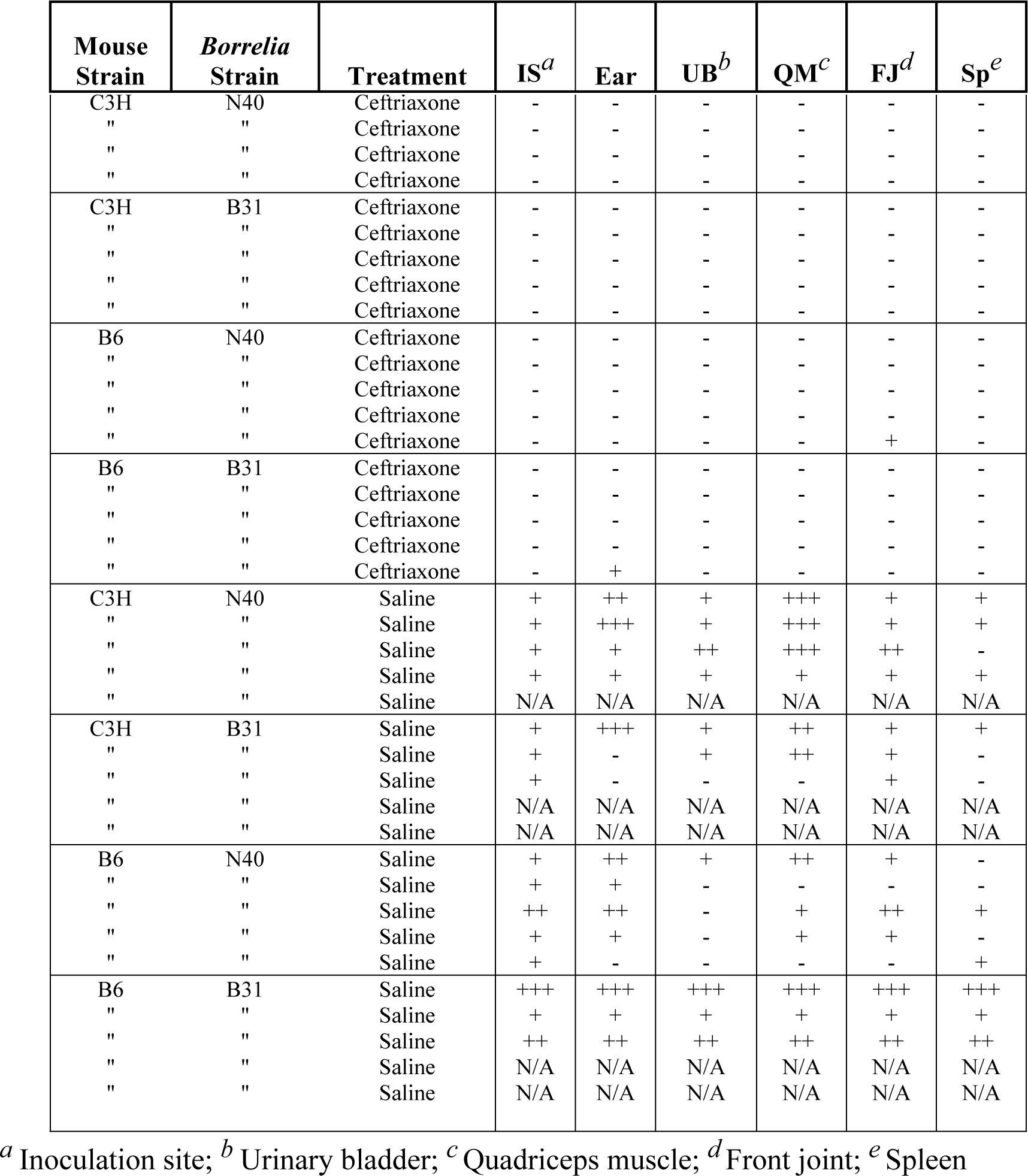
Culture results for individual mice treated with ceftriaxone and saline solution commencing at 4 weeks after inoculation, then subjected to necropsy at 18 months after completion of treatment

**Figure 3.**
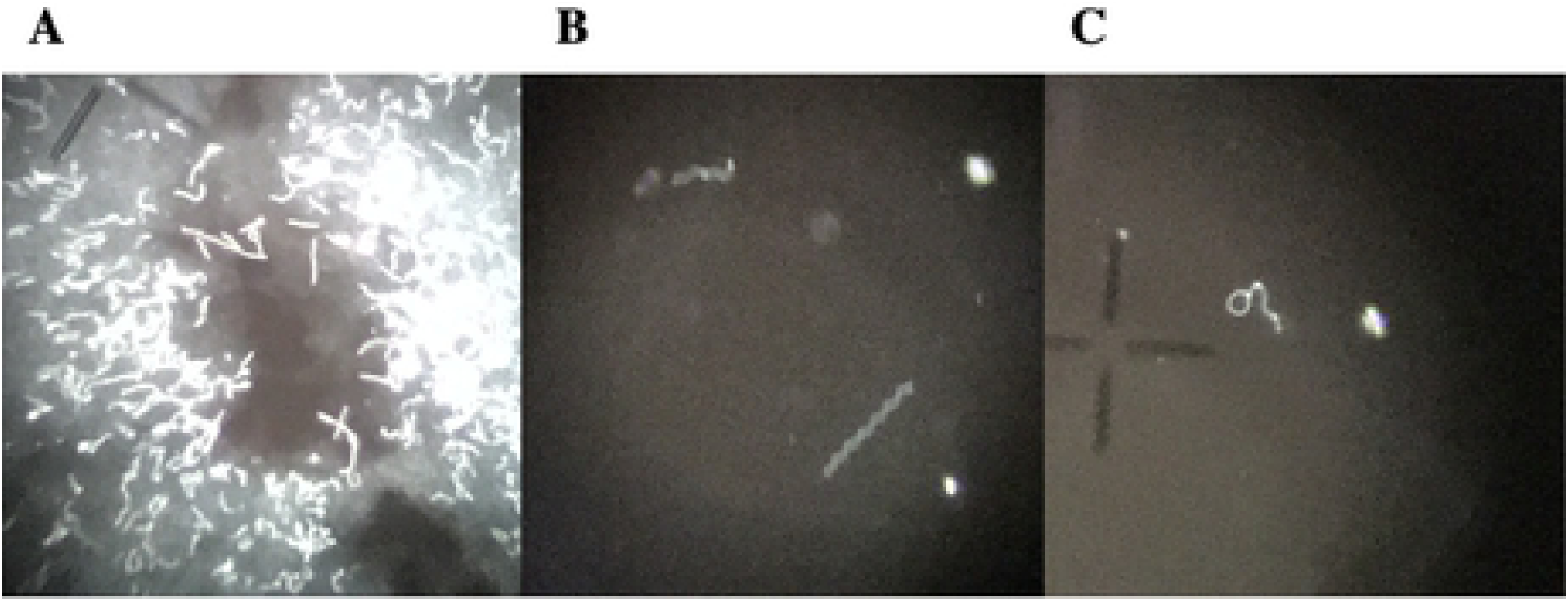
Darkfield images of cultured *B. burgdorferi* from mouse tissues necropsied 18 months after completion of treatment. Magnification 400x. **A.** Note numerous spirochetes from ear culture of saline solution treated C3H mouse infected with strain N40. **B.** Ear culture of ceftriaxone treated B6 mouse infected with strain B31. **C.** Front joint of ceftriaxone treated B6 mouse infected with strain N40

### Persistence of *B. burgdorferi* in xenodiagnostic ticks that fed upon antimicrobial-treated mice

Several days prior to necropsy, each mouse was infested by placing approximately 40 larval *I. scapularis* ticks, derived from a single population of laboratory-reared pathogen-free larvae. Ticks were allowed to feed to repletion. Engorged larvae were collected and allowed to molt into nymphs and harden, thereby allowing post-feeding amplification of spirochete populations and minimization of inhibitory products (blood) for PCR. Samples of 5 (saline treated) and 10 (ceftriaxone treated) randomly selected nymphal ticks from each tick cohort from each mouse were collected. Ticks were tested for the presence of *B. burgdorferi flaB/ospA* DNA by qPCR. Cohorts of nymphs from each mouse treated with saline solution at 12 and 18 months were all *flaB/ospA* PCR positive (Table 6). Mean copy numbers of *flaB/ospA* DNA ranged from 389 to 13,100 (mean ± SD, 3,120 ± 3,790) and from 97 to 11,950 (mean ± SD, 2,340 ± 2,820) copies per tick at 12 months and 18 months, respectively. In contrast, 9 of 20 and 3 of 19 ceftriaxone-treated mice had one *flaB/ospA* PCR positive tick each, among the ten ticks tested from each mouse at 12 and 18 months, respectively (Table 6).

**Table 6.**
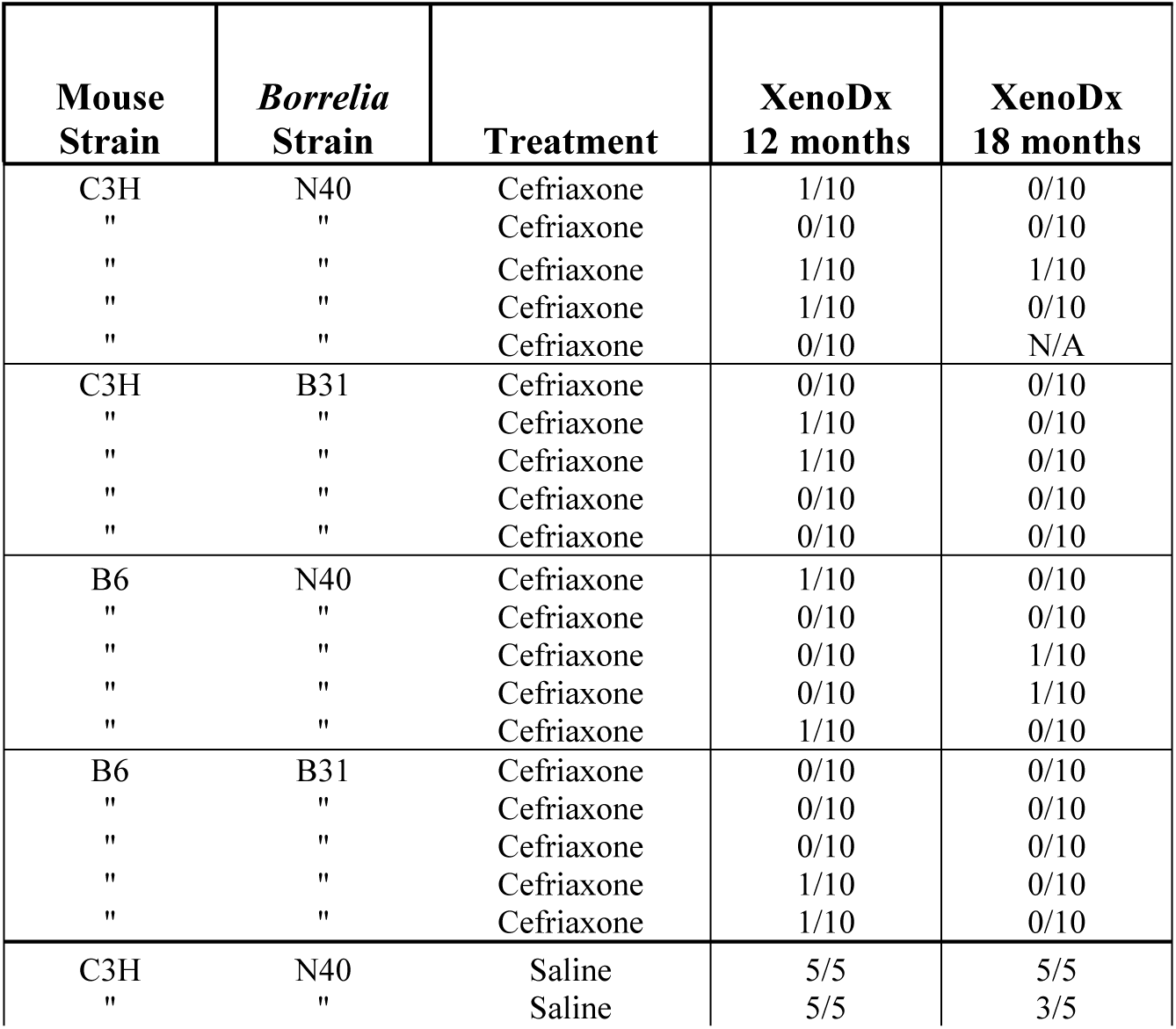
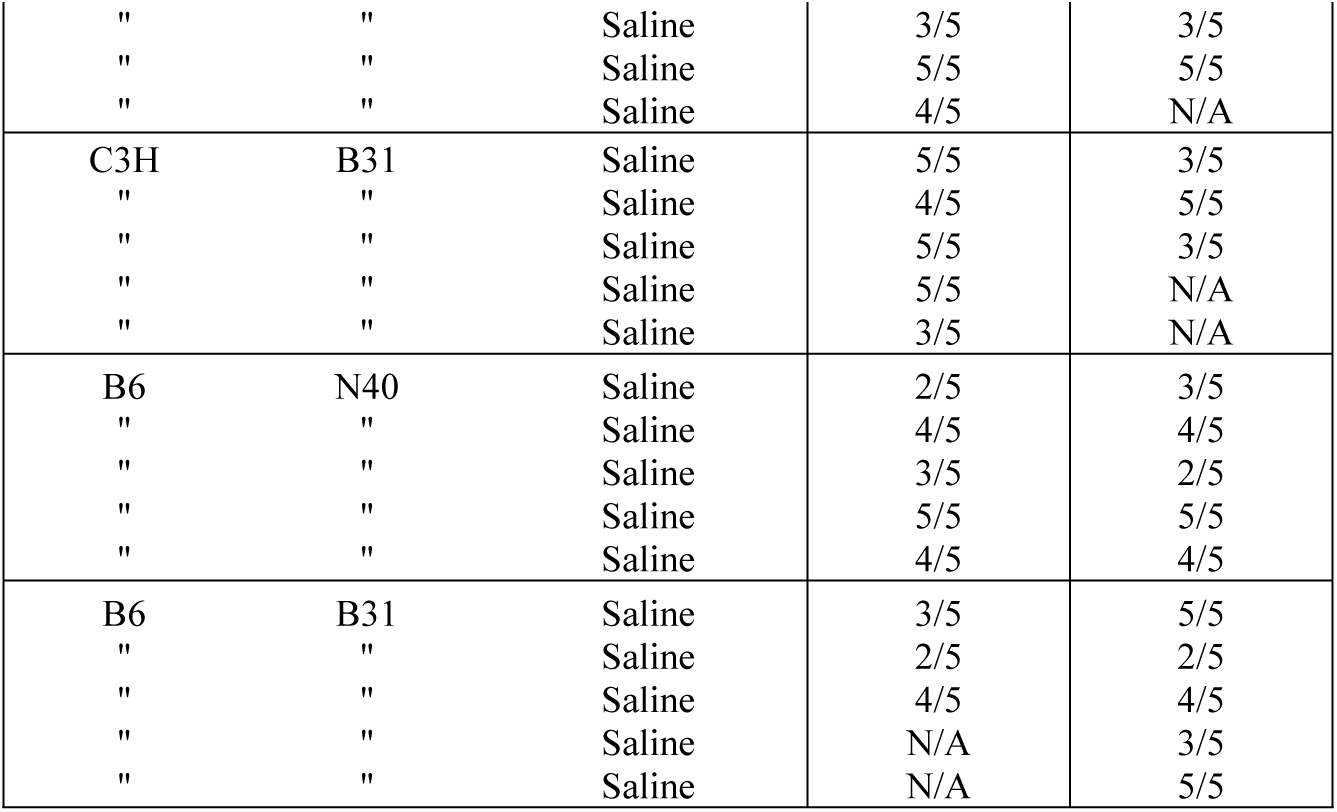
*flaB/ospA* qPCR results of nymphal that ticks fed on mice treated with ceftriaxone and saline solution prior to necropsy at 12 and 18 months after completion of treatment

### Histopathology of mouse tissues following antimicrobial treatment

Here we report on the general features of histopathology, legs were examined for evidence of arthritis and synovitis, and hearts for neutrophil and macrophage infiltration. At 12 months after treatment inflammatory scores were greatest and more robust for myocarditis in control C3H/N40 and C3H/B31 mice in comparison to antimicrobial treatment groups (Fig 4). Tibiotarsal arthritis was mild, characterized predominantly by synovial hyperplasia and hypertrophy with joint or tendon sheath effusion in both, antimicrobial treated and control mice at 12 months. There are statistically significant differences (Kruskal-Wallis test, p = 0.0227) between groups based on myocarditis scores. Group of antimicrobial C3H/B31 mice has statistically significantly lower scores than control C3H/B31 mice (Dunn’s multiple comparisons test, *p* = 0.043). No statistically significant differences (Kruskal-Wallis test, *p* = 0.1187) were observed between groups based on tibiotarsal arthritis scores. Statistical analyses performed using GraphPad Prism, 7.03 (Table 7). At 18 months inflammatory scores were greatest for myocarditis in control groups C3H/N40, C3H/B31, and B6/N40 (Fig 4). The chronic periarteritis and tenosynovitis were observed in few control C3H/B31 and B6/N40 mice, respectively, a pathohistological findings frequently observed in borrelial chronically infected mice (Fig 5). Osteoarthritis was very mild and considered a background age-related finding. There are statistically significant differences (Kruskal-Wallis test, p = 0.0019) between groups based on myocarditis scores. Control groups C3H/N40 and C3H/B31 have statistically significantly higher scores than antimicrobial treated group C3H/B31 (Dunn’s multiple comparisons test, p = 0.046, p = 0.007). No statistically significant differences (Kruskal-Wallis test, *p* = 0.1431) were observed between groups based on tibiotarsal arthritis scores. Statistical analyses were performed using GraphPad Prism, 7.03 (Table 7).

**Figure 4.**
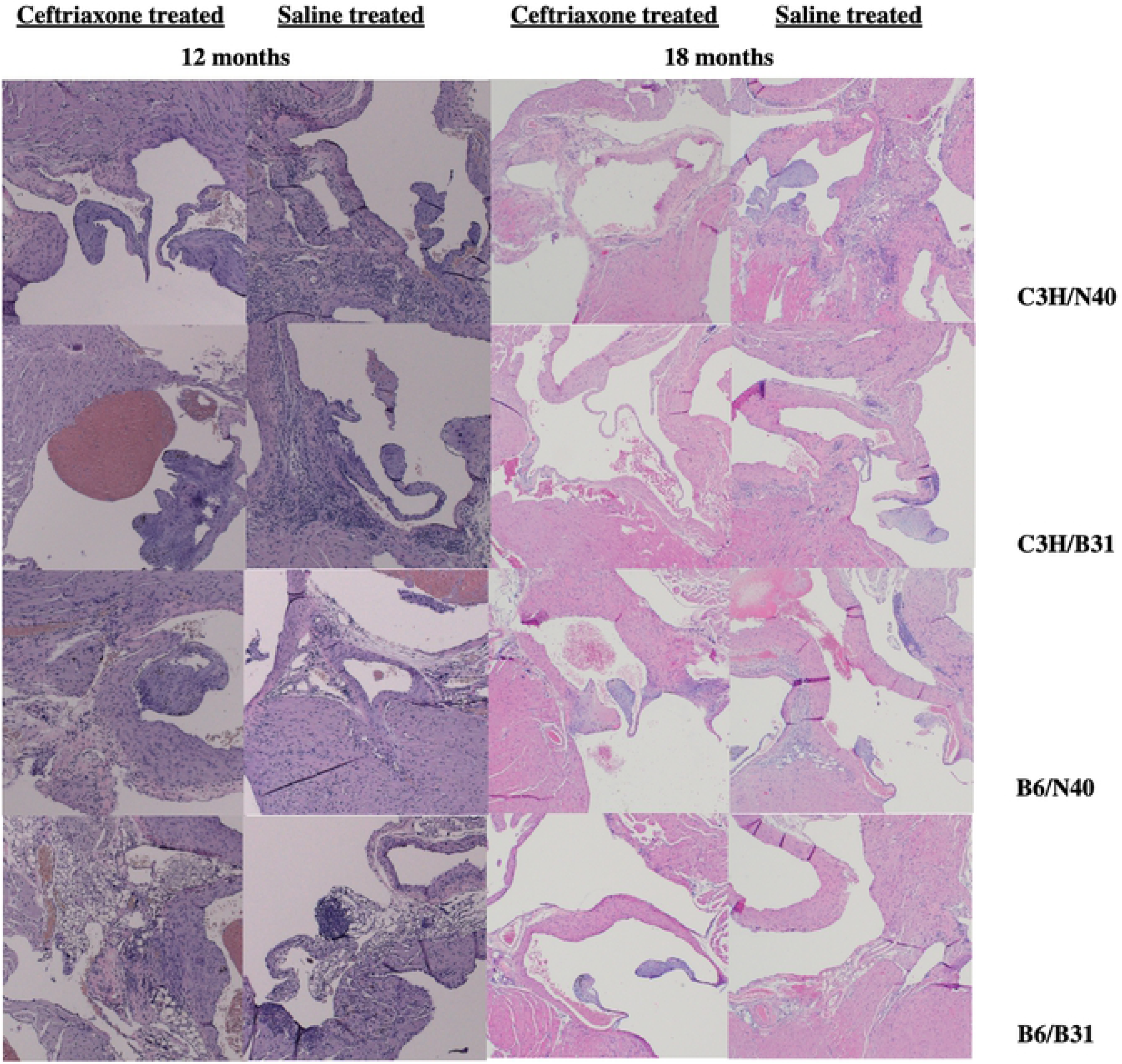
Histology of the heart base of ceftriaxone and saline solution treated mice necropsied at 12 and 18 months after treatment

**Figure 5.**
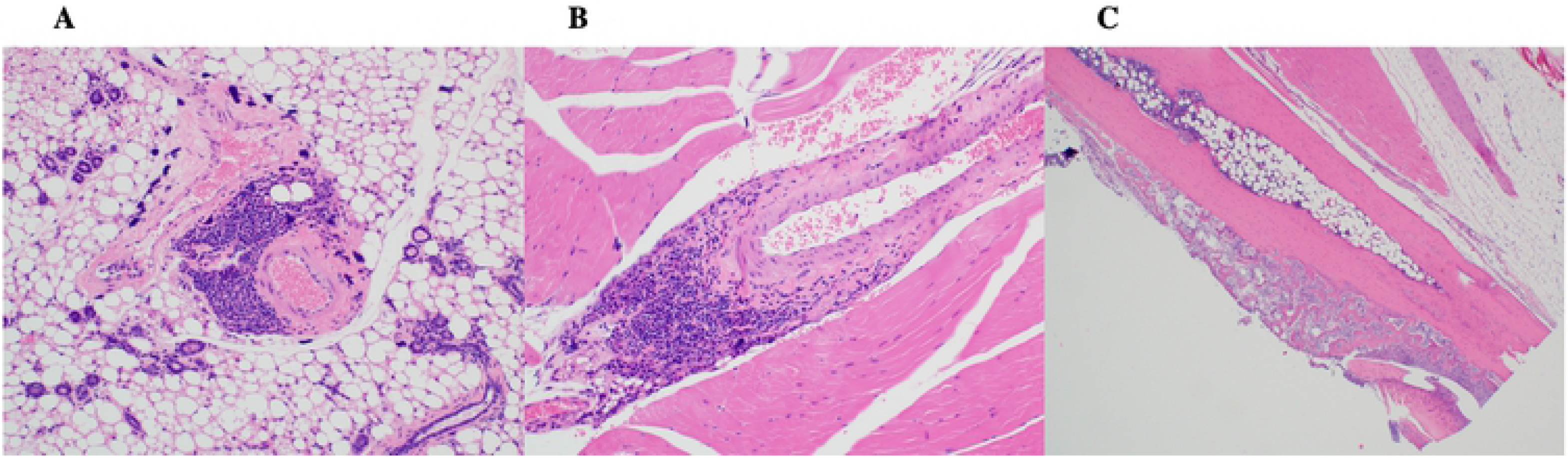
Chronic periarteritis observed in a C3H mouse infected with *B. burgdorferi* B31 then treated saline solution and necropsied 18 months after treatment (**A** and **B**). Tenosynovitis observed in a B6 mouse infected with B. burgdorferi N40 then treated with saline solution and necropsied at 18 months after treatment (**C**).

**Table 7.** Histopathological results of heart base and tibiotarsal joint of *B. burgdorferi* infected mice treated with either ceftriaxone or sham treated with saline solution

| Mouse Strain | *Borrelia* Strains | Treatment | Heart base 12 months | Tibiotarsus 12 months | Heart base 18 months | Tibiotarsus 18 months |
| --- | --- | --- | --- | --- | --- | --- |
| C3H | N40 | Ceftriaxone | 0 | 1 | 1 | 0 |
| " | " | Ceftriaxone | 0 | 0 | 1 | 0 |
| " | " | Ceftriaxone | 1 | 1 | 0 | 1 |
| " | " | Ceftriaxone | 1 | 1 | 0 | 1 |
| " | " | Ceftriaxone | 0 | 1 | N/A | N/A |
| C3H | B31 | Ceftriaxone | N/A | N/A | 0 | 0 |
| " | " | Ceftriaxone | 0 | 0 | 0 | 0 |
| " | " | Ceftriaxone | 0 | 0 | 0 | 0 |
| " | " | Ceftriaxone | 0 | 0 | 0 | 0 |
| " | " | Ceftriaxone | 0 | 0 | 0 | 0 |
| B6 | N40 | Ceftriaxone | 1 | 0 | 0 | 0 |
| " | " | Ceftriaxone | 1 | 0 | 1 | 1 |
| " | " | Ceftriaxone | 0 | 1 | 1 | 0 |
| " | " | Ceftriaxone | 1 | 0 | 0 | 1 |
| " | " | Ceftriaxone | 0 | 1 | 0 | 0 |
| B6 | B31 | Ceftriaxone | 1 | 0 | 1 | 0 |
| " | " | Ceftriaxone | 1 | 0 | 0 | 0 |
| " | " | Ceftriaxone | 0 | 0 | 1 | 0 |
| " | " | Ceftriaxone | 0 | 0 | 0 | 0 |
| " | " | Ceftriaxone | 1 | 0 | 0 | 0 |
| C3H | N40 | Saline | 1 | 0 | 1 | 1 |
| " | " | Saline | 2 | 1 | 1 | 1 |
| " | " | Saline | 1 | 1 | 2 | 1 |
| " | " | Saline | 3 | 1 | 2 | 1 |
| " | " | Saline | 1 | 1 | N/A | N/A |
| C3H | B31 | Saline | 1 | 0 | 2 | 0 |
| " | " | Saline | 1 | 1 | 2 | 0 |
| " | " | Saline | 2 | 1 | 2 | 1 |
| " | " | Saline | 3 | 1 | N/A | N/A |
| " | " | Saline | 3 | 1 | N/A | N/A |
| B6 | N40 | Saline | 0 | 1 | 1 | 0 |
| " | " | Saline | 0 | 1 | 1 | 0 |
| " | " | Saline | 1 | 0 | 1 | 1 |
| " | " | Saline | 1 | 1 | 2 | 1 |
| " | " | Saline | 1 | 1 | 1 | 1 |
| B6 | B31 | Saline | 2 | 1 | 1 | 0 |
| " | " | Saline | 1 | 1 | 0 | 0 |
| " | " | Saline | 1 | 0 | 1 | 1 |
| " | " | Saline | N/A | N/A | 1 | 0 |
| " | " | Saline | N/A | N/A | 1 | 0 |

Myocarditis scores by graduation (0-3): 0 = not present; 1 = minimal to mild, focal to focally extensive, lymphoplasmacytic infiltrates; 2 = moderate, multifocal, lymphoplasmacytic infiltrates; 3 = marked, coalescing to diffuse, lymphoplasmacytic infiltrates. Arthritis scores by graduation (0-1): 0 = not present; 1 = synovial hyperplasia/hypertrophy and effusion with or without minimal lymphoplasmacytic infiltrates

### Plasmid content of attenuated spirochetes

In our previous work we conducted experiments utilizing primarily strain N40, so the designed qPCR assays were based on its genome sequences. For analysis of plasmid presence in tissue samples from C3H and B6 mice infected with *B. burgdorferi* B31 strain, fewer genes were targeted. A comparison of plasmid profile, size, and GC content between N40 and B31 strains appear to have major structural differences, so only four designed qPCR assays targeting *bptA, arp, ospA*, and *erp23* are homologous.

Among 12-month and 18-month samples, all gene targets were detected in tissue samples from saline-treated C3H and B6 mice infected with either *B. burgdorferi* N40 and B31 strains (Table 8 and 9). *B. burgdorferi* genomes of N40 strain in tissue samples with high Bb DNA copy numbers from 12 antimicrobial-treated mice were uniformly missing *bptA* (lp25), *erp22* (cp32-9), and *erp23* (cp34-2), and variably missing *arp* (lp28-5), *eppA* (cp9), and *ospC* (cp26), whereas *ospA* (lp54) was detected in all examined tissue samples (Table 8). All assessed tissue samples of mice infected with strain B31 were uniformly missing *bptA* (lp25) and *arp* (lp28-5), and variably missing *erp23* (cp34-7). As in tissue samples from mice infected with strain N40, *ospA* (lp54) was detected in all examined samples (Table 9). Data suggest loss of small linear and circular plasmids in persisting spirochetes at 12 and 18 months after treatment with antimicrobial. No difference was observed between different *B. burgdorferi* strains (B31 and N40) or in genetically different strains of mice that are disease-susceptible (C3H) and disease-resistant (B6). Most notable was the uniform absence of lp25 in both *B. burgdorferi* strains, N40 and B31, loss of which significantly attenuates infectivity in mice, with markedly reduced, but detectable levels of infection. Results also demonstrated amplification of multiple gene targets, thereby verifying the specificity of residual BbDNA results (Tables 8 and 9).

**Table 8.** Analysis of plasmid presence in heart base (HB), ventricular muscle (VM), and tibiotarsus (TT) of mice infected with B. burgdorferi **N40** strain then treated with either ceftriaxone or sham treated with saline solution

| Mouse Strain | Treatment /Time | Sample Type | chrom *flaB* | lp25 *bptA* | lp28-5 *arp* | lp54 *ospA* | cp9 *eppA* | cp26 *ospC* | cp32-9 *erp22* | cp34-2 *erp23* |
| --- | --- | --- | --- | --- | --- | --- | --- | --- | --- | --- |
| C3H | Saline/12 | HB | + | + | + | + | + | + | + | + |
| B6 | " | TT | + | + | + | + | + | + | + | + |
| C3H | Saline/18 | HB | + | + | + | + | + | + | + | + |
| B6 | " | VM | + | + | + | + | + | + | + | + |
| C3H | Ceftriaxone /12 | HB | + | - | - | + | - | - | - | - |
| C3H | " | TT | + | - | - | + | - | + | - | - |
| C3H | " | TT | + | - | - | + | - | - | - | - |
| B6 | " | TT | + | - | + | + | - | - | - | - |
| B6 | " | HB | + | - | - | + | - | - | - | - |
| B6 | " | TT | + | - | - | + | - | - | - | - |
| C3H | Ceftriaxone<br>/18 | TT | + | - | - | + | - | - | - | - |
| C3H | " | TT | + | - | - | + | - | - | - | - |
| C3H | " | TT | + | - | + | + | + | + | - | - |
| B6 | " | TT | + | - | - | + | - | - | - | - |
| B6 | " | TT | + | - | - | + | - | - | - | - |
| B6 | " | TT | + | - | - | + | - | - | - | - |

**Table 9.** Analysis of plasmid presence in heart base (HB), ventricular muscle (VM), and tibiotarsus (TT) of mice infected with *B. burgdorferi* **B31** strain then treated with ceftriaxone and sham treated with saline solution

| Mouse Strain | Treatment/Time | Sample Type | chrom *flaB* | lp25 *bptA* | lp28-1 *arp* | lp54 *ospA* | cp34-7 *erp23* |
| --- | --- | --- | --- | --- | --- | --- | --- |
| C3H | Saline/12 | VM | + | + | + | + | + |
| B6 | " | VM | + | + | + | + | + |
| C3H | Saline/18 | HB | + | + | + | + | + |
| B6 | " | HB | + | + | + | + | + |
| C3H | Ceftriaxone/12 | TT | + | - | - | + | - |
| C3H | " | HB | + | - | - | + | - |
| C3H | " | TT | + | - | - | + | - |
| B6 | " | TT | + | - | - | + | + |
| B6 | " | HB | + | - | - | + | - |
| B6 | " | TT | + | - | - | + | - |
| C3H | Ceftriaxone/18 | TT | + | - | - | + | - |
| C3H | " | TT | + | - | - | + | - |
| C3H | " | TT | + | - | - | + | - |
| B6 | " | TT | + | - | - | + | - |
| B6 | " | TT | + | - | - | + | - |
| B6 | " | TT | + | - | - | + | - |

## DISCUSSION

The most commonly prescribed antimicrobial agents for the treatment of human Lyme borreliosis, such as penicillin, amoxicillin, cefotaxime, cefuroxime, ceftriaxone, doxycycline, and erythromycin, have shown to be effective against *B. burgdorferi* (72, 73). It is important to mention that treatment during early infection is more effective in clearing the pathogen than treatment during later infection in humans (74) and in the mouse model (46). Over the past several years, there have been numerous well-supported reports of *Borrelia* detection after completed antimicrobial therapy (46, 50, 53, 60, 75, 76). All of these studies have in common that spirochetes were detected by PCR for BbDNA, but not by culture. Morphologically intact spirochetes were visualized by immunohistochemistry in tissues from antimicrobial-treated mice; ticks could acquire those spirochetes and transmit them to recipient mice. Mice tissues were positive for *B. burgdorferi*-specific RNA transcripts.

In the current study, we evaluated persistence at 12 and 18 months after ceftriaxone treatment in two strains of mice that are disease-susceptible (C3H) and disease-resistant (B6). We recognize that inbred mice do not represent genetically heterogeneous humans but use of two genetically and phenotypically disparate mouse strains will confirm the generality of persistence, and more importantly resurgence, following antimicrobial treatment. Disease severity and tissue spirochete burdens vary among mouse strains (77), although the ID50 among mice is identical (78). C3H/HeJ mice are susceptible to infection by Gram-negative bacteria as a result of spontaneous mutation in *Tlr4* gene (79), which may influence the persistence. B6 mice harbor equivalent tissue spirochete burdens as C3H mice, become persistently infected, and have normal macrophage function (79).

Our results have shown that, indeed, persistence and resurgence in fully immunocompetent B6 mice is similar to those in disease-susceptible C3H mice, which confirm the generality of persistence in a mouse model. These results confirm our previous finding (47) that a subpopulation of viable, antimicrobial-tolerant, but slowly dividing, and persistent spirochetes of *B. burgdorferi* resurged in mice 12 months after treatment and re-disseminated into multiple tissues. Importantly, copy numbers of spirochetal DNA in tissue samples of antimicrobial treated disease-resistant B6 mice was nearly equivalent to those of disease-susceptible C3H mice. However, significantly higher DNA copy numbers of spirochetal DNA were determined in saline-treated groups of C3H mice infected with either N40 or B31 in comparison to antimicrobial-treated mice (C3H and B6). This finding is in concordance with the results published by Armstrong et al. (80) that C3H mice develop more severe disease than B6 mice, with greater spirochetes number and later clearance.

Despite the fact that tissues of treated mice remained BbDNA PCR-positive at 12 months with significant copy numbers of targeted genes, culture was consistently negative. Actually, in a single cultured urinary bladder we observed two non-motile spirochetes for up to three weeks of incubation. On the other hand, all saline-treated mice were culture-positive. Our studies (46, 47, 50) and those of others (45, 49, 81) in mice, dogs (29, 52) and non-human primates (53, 54) have all reached similar conclusions: spirochetes are persisting, but are paradoxically non-cultivable. It was suggested that the persisting remnants of *B. burgdorferi* in the tissues of infected mice after antimicrobial treatment is DNA or DNA-containing structures rather than live bacteria (76, 82, 83). In their *in vitro* study Iyer et al. (84) could not successfully subculture spirochetes after exposure to ceftriaxone. However, BbDNA was detected by PCR for up to 56 days in aliquots from both ceftriaxone-treated and untreated cultures. Pavia and Wormser (85) demonstrated that *B. burgdorferi* cannot be cultured from experimentally infected mice after ceftriaxone treatment with only 5 daily doses. The treatment regimen used in the study was judged based on the absence of a positive culture.

IDSA Guidelines state that “unless proven otherwise, culture should be regarded as the gold-standard to address viability of *B. burgdorferi*”(86). Culture may indeed be a gold standard when it is positive, but it is often not. It is apparent that not all isolates or strains can be easily cultured, and this is especially apparent during long-term infection. It is becoming increasingly clear that in *B. burgdorferi* infected and antimicrobial treated animals, spirochetes were non-cultivable but viable, as have been demonstrated by xenodiagnoses (46, 47, 50, 54), immunohistology (46, 47, 75), and transcriptional activity of mRNA (47, 50, 54). Here we demonstrated that morphologically intact *B. burgdorferi* that survived antimicrobial treatment in disease-susceptible C3H mice as well in disease-resistant B6 mice could be acquired by larval ticks. Ticks remained BbDNA-positive through molting into nymphs. In addition, transcription of the chromosomally-encoded *B. burgdorferi 16S rRNA* and *flaB* genes were detected in treated mice, indicating that spirochetes are metabolically active and alive at 12 months after treatment. The data obtained in this study indicate that in a significant number of culture-negative tissue samples following antimicrobial treatment, complementary methods in diagnostic microbiology should be considered. The clinical significance and the prognostic value of these findings have to be more deeply investigated. It was shown that dead bacterial DNA can be detected for up to 4 to 5 months after antimicrobial treatment (87), as extracellular DNA is very prone to degradation (88, 89). Bacterial mRNAs, on the other hand, have very short half-lives as degraded by exonucleases very fast. The half-life ranges from a few minutes to several hours, depending on the bacterial strain (90-92). These finding support the fact that non-cultivability of antimicrobial-tolerant and persistent spirochetes does not negate their viability.

Our previous studies in mice (46, 47, 50) and those of others (49, 81) are all based on *B. burgdorferi* N40 strain. Although N40 probably represents *B. burgdorferi*, the generality of long-term spirochete persistence following antimicrobial treatment should be evaluated with other *B. burgdorferi* strains. Therefore, we used *B. burgdorferi* strain B31 and confirmed persistence and resurgence at 12 months after antimicrobial treatment in both, C3H and B6, mice. Our results indicate that disseminated spirochetes of two different *B. burgdorferi* strains can persist in mice at 12 and 18 months following antimicrobial treatment. Non-cultivable spirochetes persist in mice (93), dogs (29), and non-human primates (54) inoculated with alternate strains of *B. burgdorferi*, so we are confirming that persistence and resurgence are not unique to N40.

An additional long-term interval, holding mice up to 18 months, facilitated several observations regarding persistent antimicrobial-tolerant spirochetes. Within the observation period, there was molecular evidence of BbDNA in 79% of all assessed mice following antimicrobial treatment. Spirochetal DNA was exclusively detected in tibiotarsal joint, but not in heart base and ventricular muscle. Copy numbers of BbDNA are not significantly different between treated C3H and B6 mice infected with either N40 or B31. Interestingly, BbDNA copy numbers in tibiotarsal joints of all antimicrobial treated mice at 18 months are equivalent to those at 12 months, suggesting that antimicrobial-tolerant spirochetes remained persistent exclusively in connective-tissue. One of the proposed mechanisms of *B. burgdorferi* immune evasion and persistence is sequestration in connective-tissue, that make them less accessible to cells and molecules of the host’s immune system (55). It was suggested that the joint or a tissue adjacent to the joint is the niche of persisting *B. burgdorferi* in antimicrobial-treated mice (49). There is scientific evidence that tissues with greater decorin expression levels such as joint harbored most spirochete loads during chronic infection (94). The findings of our study reinforce the notion of persistence of viable spirochetes following antimicrobial treatment and further implicate the connective-tissue as a privileged site that may partake to its failure. Although spirochetes resurged 12 months after antimicrobial treatment, overt disease was not present. Also, despite spirochetes presence, postmortem gross signs of disease were not observed in assessed tissues (heart, joint) of treated mice at 18 months. Studies in animal models have shown that resolution of arthritis and carditis is mediated by the acquired humoral immune response of the host. Under these conditions, anatomically defined inflammation resolves, but infection persists (95-97). Indeed, even during the pre-immune phase of infection, spirochetes populate many tissues with no evidence of inflammation (thus inflammation does not necessarily correlate with spirochete presence).

Spirochetal RNA was detected in the joint tissue of C3H and B6 mice infected with either N40 or B6 following treatment at 18 months, suggesting their viability. Interestingly, xenodiagnosis was positive in only three of nineteen treated animals. We are speculating that the inability to retrieve spirochetes from all treated mice could be ascribed to a clearance of spirochetes from the heart tissue. Migration of spirochetes from joints to tick attachment sites, that is usually a head and/or a dorsal part of a mouse body, is more distant than from a heart. It was also suggested that the efficiency of spirochetal acquisition from a host to a tick depend on the intensity and duration of infection, and it is significantly less during chronic infection (98). However, we recovered a few slow-growing spirochetes by culture from joint tissues of two mice, utilizing several specially prepared media that were cordially provided by Monica E. Embers. We were unable to propagate the recovered spirochetes as they lysed after 4 weeks. Obviously, the used media were unable to support the prolonged growth of slow growing and impaired spirochetes.

Here we have shown that following antimicrobial treatment, both strains of *B. burgdorferi*, N40 and B31, generates an increasingly heterogeneous population of replicatively-attenuated spirochetes that have lost one or more small plasmids. These “attenuated” spirochetes remain viable, but because of their plasmid loss, they divide slowly, thereby being tolerant to the effects of antimicrobials, as well as being non-cultivable. The phenomenon of persistent, non-cultivable *B. burgdorferi* spirochetes after antimicrobial treatment has been explained by antimicrobial tolerance. Unlike antimicrobial-resistance, antimicrobial-tolerance to all classes of antimicrobials fail to completely eliminate non-dividing or slowly-dividing subpopulations of a broad array of bacteria and fungi (62, 99). It has been shown that *in vitro* passage of *B. burgdorferi* is predisposed to plasmid loss (63-65). This phenomenon probably occurs throughout the infection and elevates over time.

Clinical relevance of microbial persistence has been demonstrated for *Burkholderia pseudomallei* (100), *Escherichia coli* (101), *Pseudomonas aeruginosa* (102), *Candida albicans* (103), *Streptococcus pneumonia* (104), *Staphylococcus aureus* (101, 105), *Mycobacterium tuberculosis* (106, 107), *Legionella* spp. (108, 109), and *Salmonella enterica* (110). The persistent cells have transient tolerance to antimicrobial agents and their ability to revert back to a sensitive state, in which cells rapidly divide make them important in chronic infections. There are two major consequences of the presence of persister cells: 1) the continuous presence of viable cells during consecutive rounds of antimicrobial treatment, which contribute to the emergence of antimicrobial resistance and 2) experimental evidence indicates that prolonged exposure to antimicrobial agents leads to the selection of high persistent mutants (111, 112). The biological relevance of attenuated *B. burgdorferi* spirochetes is probably inconsequential, while their clinical relevance was subject of this study. Our study demonstrated that non-cultivable spirochetes can persist in a host following antimicrobial treatment for a long time, but did not demonstrate their clinical relevance in a mouse model of chronic infection. The clinical relevance of attenuated spirochetes following antimicrobial treatment and their eventual fate in other animal models require further studies.

## ACKNOWLEDGMENT

The technical assistance of Kim Olsen and Cara Wademan was essential to this study and is greatly appreciated. We thank Dr. Melissa J. Caimano for providing *I. scapularis* larval ticks and Dr. Monica E. Embers for providing culture media.

## AUTHORS CONTRIBUTION

Conceived and designed the experiments: EH. Performed the experiments: EH DI EE. Analyzed the data: EH DI EE. Contributed reagents/materials/analysis tools: DI EE. Wrote the paper: EH DI.

